# Reduction of *Paraoxonase* expression followed by inactivation across independent semiaquatic mammals suggests stepwise path to pseudogenization

**DOI:** 10.1101/2022.09.29.510191

**Authors:** Allie M Graham, Jerrica M Jamison, Marisol Bustos, Charlotte Cournoyer, Alexa Michaels, Jason S Presnell, Rebecca Richter, Daniel E. Crocker, Ari Fustukjian, Margaret E. Hunter, Lorrie D. Rea, Judit Marsillach, Clement E Furlong, Wynn K Meyer, Nathan L Clark

**Affiliations:** Department of Human Genetics, University of Utah, Salt Lake City, UT; Department of Biological Sciences, University of Toronto - Scarborough, Scarborough, Ontario, Canada; Department of Biomedical Engineering, University of Texas – San Antonio, San Antonio, TX; South Florida Wildlife Center, Fort Lauderdale, FL; Graduate School of Biomedical Sciences, Tufts University, Boston, MA; The Jackson Laboratory, Bar Harbor, ME; Department of Medicine, Division of Medical Genetics, University of Washington, Seattle, WA; Department of Biology, Sonoma State University, Rohnert Park, CA; Loveland Living Planet Aquarium, Draper, UT; U.S. Geological Survey, Wetland and Aquatic Research Center, Gainesville FL, USA, 32653; Water and Environmental Research Center, Institute of Northern Engineering, University of Alaska – Fairbanks, Fairbanks, AK; Department of Environmental & Occupational Health Sciences, University of Washington School of Public Health, Seattle, WA; Department of Genome Sciences, University of Washington, Seattle, WA; Department of Biological Sciences, Lehigh University, Bethlehem, PA

**Author notes:** contributed equally (Wynn K Meyer, Nathan L Clark). Corresponding authors: Nathan Clark Wynn Meyer Allie Graham.

## Abstract

Convergent adaptation to the same environment by multiple lineages frequently involves rapid evolutionary change at the same genes, implicating these genes as important for environmental adaptation. Such adaptive molecular changes may yield either change or loss of protein function; loss of function can eliminate newly deleterious proteins or reduce energy necessary for protein production. We previously found a striking case of recurrent pseudogenization of the *Paraoxonase 1* (*Pon1*) gene among aquatic mammal lineages - *Pon1* became a pseudogene with genetic lesions, such as stop codons and frameshifts, at least four times independently in aquatic and semiaquatic mammals. Here, we assess the landscape and pace of pseudogenization by studying *Pon1* sequences, expression levels, and enzymatic activity across four aquatic and semiaquatic mammal lineages: pinnipeds, cetaceans, otters, and beavers. We observe in beavers and pinnipeds an unexpected reduction in expression of *Pon3*, a paralog with similar expression patterns but different substrate preferences. Ultimately, in all lineages with aquatic/semiaquatic members, we find that preceding any coding level pseudogenization events in *Pon1*, there is a drastic decrease in expression, followed by relaxed selection, thus allowing accumulation of disrupting mutations. The recurrent loss of *Pon1* function in aquatic/semiaquatic lineages is consistent with a benefit to *Pon1* functional loss in aquatic environments. Accordingly, we examine diving and dietary traits across pinniped species as potential driving forces of *Pon1* functional loss. We find that loss is best correlated with diving activity and likely results from changes in selective pressures associated with hypoxia and hypoxia-induced inflammation.

## INTRODUCTION

Gene loss and pseudogenization events occur regularly over the course of evolution. Genes contributing to evolutionary fitness are rarely lost due to selection against such loss, but genes under relaxed constraint, or no constraint at all, are frequently lost during evolution. For instance, there are multiple cases in which certain genes appear to become expendable as a species moves into a new environment, such as the large-scale loss of function of genes mediating olfaction, as well as both deletion and loss of function of taste receptor genes in marine mammals (Hayden, et al. 2010; Feng, et al. 2014; Zhu, et al. 2014; Chikina, et al. 2016). However, recent work has suggested that gene functional loss may serve as an engine of evolutionary change (Olson 1999b), and it has been implicated in the evolution of lineage-specific traits presumably crucial for fitness (Zhao, et al. 2010; Meredith, et al. 2011; Albalat and Cañestro 2016; Yohe, et al. 2017; Esfeld, et al. 2018; Graham and Barreto 2020). For example, the pseudogenization of tooth gene enamelysin in certain cetaceans has been suggested to have led to the formation of baleen (Meredith, et al. 2011), functional loss of the *T1R1* umami taste receptor gene in giant panda is associated with its switch to bamboo consumption (Zhao, et al. 2010), and a variety of animals have lost function in short-wave opsins in a wide array of light-limited environments (Carvalho, et al. 2006; Zhao, et al. 2009; Davies, et al. 2012; Weadick, et al. 2012; Cortesi, et al. 2015; EscobarϑCamacho, et al. 2017).

In many such cases, gene loss (or pseudogenization) could have been preferentially selected due to the adaptive advantage such functional loss provides. Evidence for adaptive gene loss comes in many forms. When different species encounter the same selective pressure, their convergent loss of a particular gene could support the hypothesis of adaptive evolution. More generally, it has been argued that the observation of convergence in any trait, including convergent loss of function, is in itself evidence that the trait is adaptive (Gompel and Prud’homme 2009; Losos 2011). However, convergent gene loss could also arise due to convergent relaxation of constraint on a gene (Lahti, et al. 2009). There are several examples of independent losses or pseudogenization events of the same gene, belying convergent selective pressures (McGrath, et al. 2011; Davies, et al. 2012; Policarpo, et al. 2021). Other clues to the type of selective shift underlying gene loss could come from the rate at which a gene is lost, with rapid loss suggesting an adaptive advantage. Sharma et al., for instance, argue that loss that is contemporaneous with a particular phenotypic adaptation provides evidence for the contribution of the gene loss to the adaptation (Sharma, et al. 2018). In contrast, gene loss due to relaxation of constraint would be expected to follow the phenotypic adaptation. Additional evidence of adaptive gene loss could be the presence of population genetic signatures of positive selection associated with the gene locus.

An example of convergent gene functional loss that was likely adaptive, with evidence of an advantage to gene deactivation, is seen in the deterioration of ocular genes in cave fish. In a nutrient-poor cave environment, genetic lesions that led to eye regression are thought to have been advantageous because the eye consumes a large amount of energy (Moran, et al. 2015). In that case, genetic changes leading to regressed eyes were initially adaptive, but subsequent gene losses could likely be due to relaxed constraint (Jeffery 2005; Yoshizawa, et al. 2012; Cartwright, et al. 2017). There are multiple additional examples of pseudogenization associated with adaptive traits, most notably eye-loss in other cave-dwelling animals (Fang, et al. 2014; Moran, et al. 2015), coloration in birds (Borges, et al. 2015; Emerling 2018), diet (Hecker, et al. 2019), olfaction in Drosophila (Dekker, et al. 2006; Prieto-Godino, et al. 2016), host-pathogen interactions (Sun, et al. 2008; Key, et al. 2014), teeth/dentition (Meredith, et al. 2011; Meredith, et al. 2014), and lactation/placentation (Brawand, et al. 2008).

We previously found a striking case of recurrent pseudogenization of the *Paraoxonase 1* (*Pon1*) gene among aquatic and semiaquatic mammal lineages (Meyer, et al. 2018). *Pon1* became a pseudogene at least four times in aquatic and semiaquatic mammals with clear genetic lesions, such as stop codons and frameshifts. *Pon1* function was lost in the ancestor of all modern cetaceans (whales, dolphins), in the ancestor of sirenians (manatees, dugongs), within pinnipeds (seals, sea lions), and in sea otters. In contrast, no genetic lesions in *Pon1* have been found in any of the 59 terrestrial mammal species we have examined to date.

The question remains as to whether the loss of *Pon1* function in aquatic and semiaquatic lineages was facilitated by directional selection because pseudogenization was adaptive, or if it is simply the result of convergent relaxed constraint. Colonizing aquatic niches involves exposure to challenging environmental and selective pressures, resulting in a range of convergent adaptations influencing aspects of these species’ biology from physiology and morphology to the genome(Fish, et al. 2008; Chikina, et al. 2016; Sharma, et al. 2018; Davis 2019). In humans, *PON1* is expressed predominantly in the liver; it encodes a bloodstream enzyme that reduces oxidative damage to lipids in low-and high-density lipoprotein (LDL and HDL) particles and is associated with the prevention of atherosclerotic plaque formation (Mackness, et al. 1991; Rosenblat and Aviram 2009; Précourt, et al. 2011; Aharoni, et al. 2013). PON1 also hydrolyzes the oxon forms of specific organophosphate compounds, such that it is the main line of defense against some man-made pesticide by-products including chlorpyrifos oxon and diazoxon (Li, et al. 2000; Costa, et al. 2005). Thus, it was hypothesized that *Pon1*’s functional loss in marine mammals was potentially related to lipid metabolism and/or oxidative stress response encountered from repeated diving (Furlong, et al. 2016; Meyer, et al. 2018). This is because aquatic mammal diets differ in the types and proportion of certain polyunsaturated fatty acids (PUFAs), which show differing levels of oxidative damage (Miyashita, et al. 1993; Koussoroplis, et al. 2008). Another hypothesized reason for this convergence on *Pon1* functional loss also involves oxidative damage – breath-hold diving is associated with transient hyperoxia followed by hypoxia and a build-up of carbon dioxide, chest-wall compression and significant hemodynamic changes (Zenteno-Savın, et al. 2012).

Here, we assess the landscape and pace of *Pon1* pseudogenization by assessing sequence data, expression levels, and biochemical activity across two fully aquatic mammalian clades (pinnipeds and cetaceans) and two clades with semiaquatic members (beavers within rodents, otters within mustelids). In addition, we examine potential driving forces of *Pon1* loss through correlation tests with behavioral and physiological traits related to dive time and diet. Ultimately, in all lineages with aquatic/semiaquatic members, we find that preceding any coding level pseudogenization events in *Pon1*, there is evidence of significant decrease in expression, followed by relaxed selection, thus allowing for an accumulation of disrupting mutations. We suggest that loss of *Pon1* is best correlated with diving activity, and likely the result of similar pressures associated with hypoxia and hypoxia-induced inflammation processes.

## RESULTS

We took a multi-faceted approach to determine the extent and timing of *Pon1* functional losses and to assess evidence for loss of function at DNA, RNA, and protein levels in aquatic and semiaquatic mammal groups (Figure 1). For a variety of new species selected for aquatic/non-aquatic contrasts, we identified or generated *Pon1* coding sequences, collected data for liver mRNA levels of *Pon1* and *Pon3*, and measured biochemical activity against Pon1 substrates in blood plasma. Based on the combined results, we infer that *Pon1* has lost function at least four times in semiaquatic mammals and that reduced expression of both *Pon1* and *Pon3* is common in these species. We present these results in subsequent sections organized around taxonomic groups for ease of interpretation.

**Figure 1:**
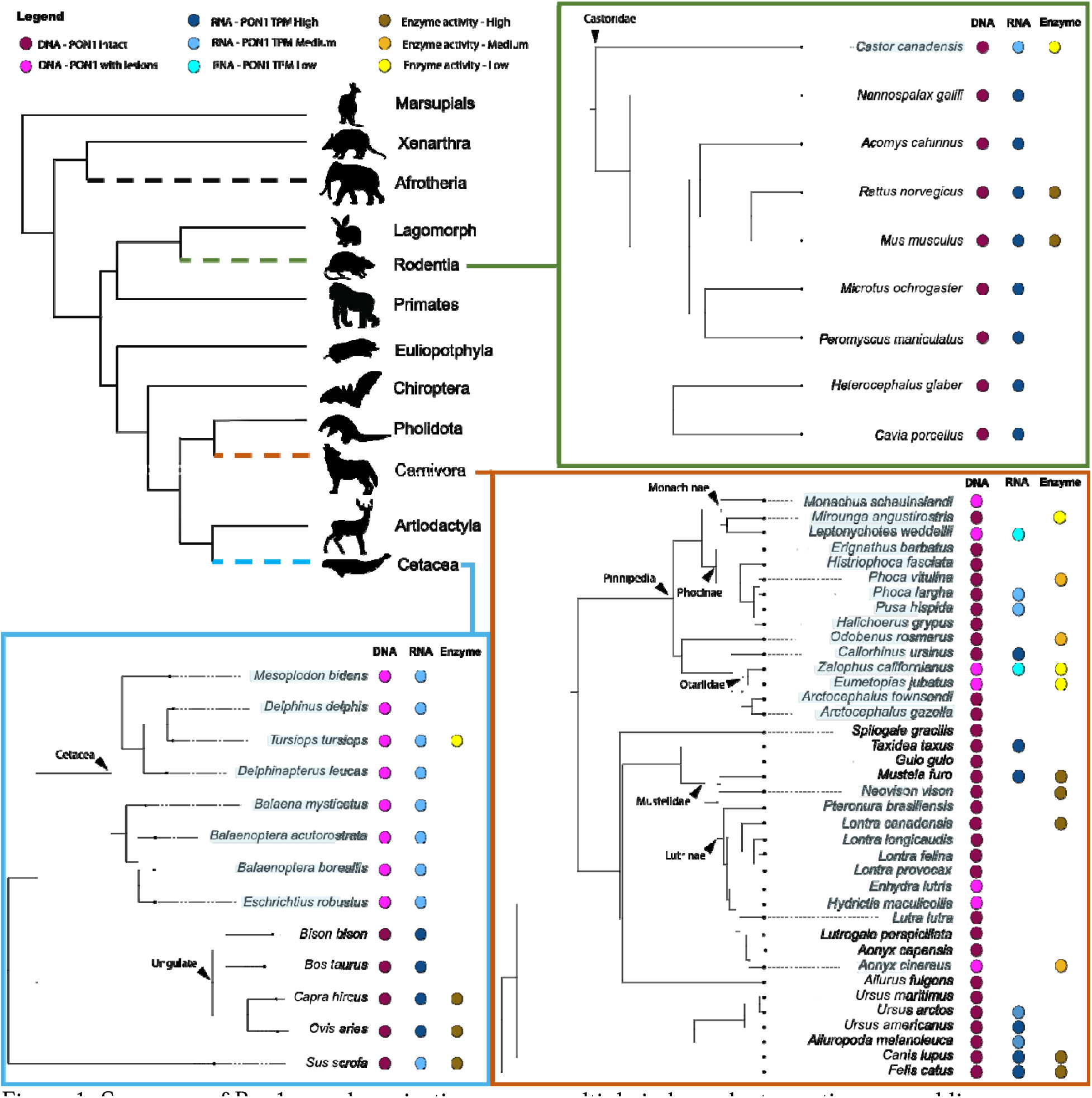
Summary of Pon1 pseudogenization across multiple independent aquatic mammal lineages - species whose labels are highlighted in light blue are aquatic or semiaquatic. The colored circles represent presence of results from DNA, RNA, or enzymatic tests; no circle means there are no data to report. Phylogenetic relationships shown are based on data from previously published work (Esselstyn, et al. 2017; Swanson, et al. 2019; McGowen, et al. 2020; de Ferran, et al. 2022).

### Pinnipeds show two instances of Pon1 functional loss and reduced Pon3 expression

We identified at least two separate pseudogenization events of *Pon1* in pinnipeds. One occurred in Otariinae (sea lions), apparently after this clade split with its sister clade approximately 5.2 Mya (Higdon, et al. 2007) (Figure 1). The California and Steller sea lions share a nonsense and a frameshift coding sequence substitution, and neither species of Arctocephalus (fur seals) in our dataset displays either these or any other inactivating mutations in their coding sequence. The blood plasma of both Otariinae sea lions also fails to metabolize organophosphate substrates or phenyl acetate (Figure 2; Supp Figure S1), confirming the loss of Pon1 function in this clade. Several species of Monachinae also display different types of evidence of Pon1 functional loss (*Leptonychotes weddellii*, *Mirounga angustirostris, and Monachus schauinslandi*), whereas the only species of the closely related Phocinae within our dataset (*Phoca vitulina*) retains some enzymatic activity, albeit reduced, against organophosphate substrates in its blood plasma (Figure 2; Supp Figure S1). This implies that the loss within Monachinae most likely occurred on the branch ancestral to all extant species within this clade, roughly 11.3 - 16 MYA (Higdon, et al. 2007). Moreover, this loss likely involved changes to a regulatory site or sites, given the lack of shared lesions across Monachinae species (Meyer, et al. 2018).

**Figure 2:**
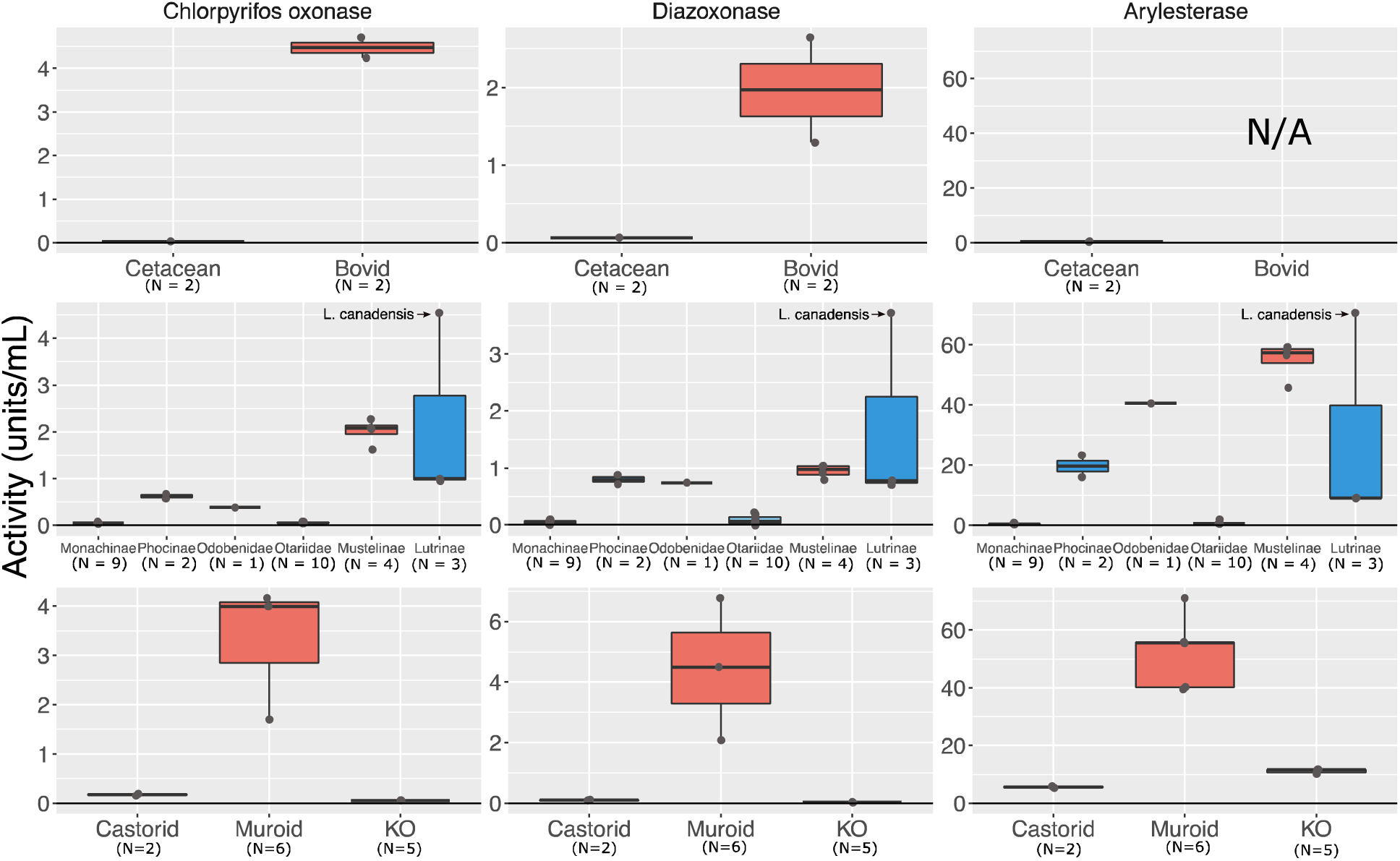
Blood plasma from aquatic and semiaquatic species varies in enzymatic activity against Pon1 substrates. Plots show enzymatic activity in units/mL against (from L to R) chlorpyrifos oxon, diazoxon, and phenyl acetate for aquatic/semiaquatic and closely related terrestrial species within (from top to bottom) Cetartiodactyla, Carnivora, and Rodentia. Carnivora species included are *M. angustirostris* (Monachinae), *P. vitulina* (Phocinae), *O. rosmarus* (Odobenidae), *E. jubatus* (9) and *Z. californianus* (1) (Otariidae), *M. furo* (Mustelinae), and *A. cinereus* (2) and *L. canadensis* (1) (Lutrinae). Data for Bovids (sheep, goat) and one Muroid are from Furlong, et al. (2000a). See also Figure S1 and Table S12.

Expression levels of *Pon1* in the liver, the sole site of *Pon1* gene expression, are typically strongly reduced in pinnipeds (Figure 3). When contrasting 5 pinniped species against 11 terrestrial carnivore species, *Pon1* expression levels are significantly lower in pinnipeds (*Mann-Whitney U test; Z =* 2.492*, P=* 0.013). The notable exception is the pinniped *Callorhinus ursinus* (Northern fur seal), which had levels within the normal range of terrestrial carnivores (Figure 3). Notably, this species has the shortest maximum dive time of all pinnipeds in this study (Figure 4; Supp Table S1). The other four pinniped species showed expression levels below those of all observed terrestrial species, and many of the pinniped expression levels were close to zero. Importantly, species in the genus *Phoca* showed reduced expression (61.76 TPM) despite having no apparent genetic lesions (*i.e.,* substitutions inducing frameshifts or stop codons that would lead to pseudogenization). This result explains the intermediate enzymatic activity in plasma for *Phoca* species (Figure 1). In contrast, expression levels of *Pon2* — a *Pon* family member expressed in nearly all tissues — and *Actin* were not different between pinnipeds and carnivores (*Pon2*, *n1=11, n2 = 5, Z = 27, P=* 1; *Actin*, *n1=11, n2 = 5, Z = 0, P= 1)*. Expression of *Pon3*, however, was also significantly and unexpectedly reduced in pinnipeds (*n1=11, n2 = 5, Z =* 2.26576, *P=* 0.0232) (Supp Table S2). However, we observed no evidence of genetic lesions in coding regions or substitutions at important sites in Pon3 that were specific to pinnipeds; the only functionally relevant site with a substitution observed in a pinniped (*O. rosmarus*) is also variable in terrestrial carnivores (Supp Table S3).

**Figure 3:**
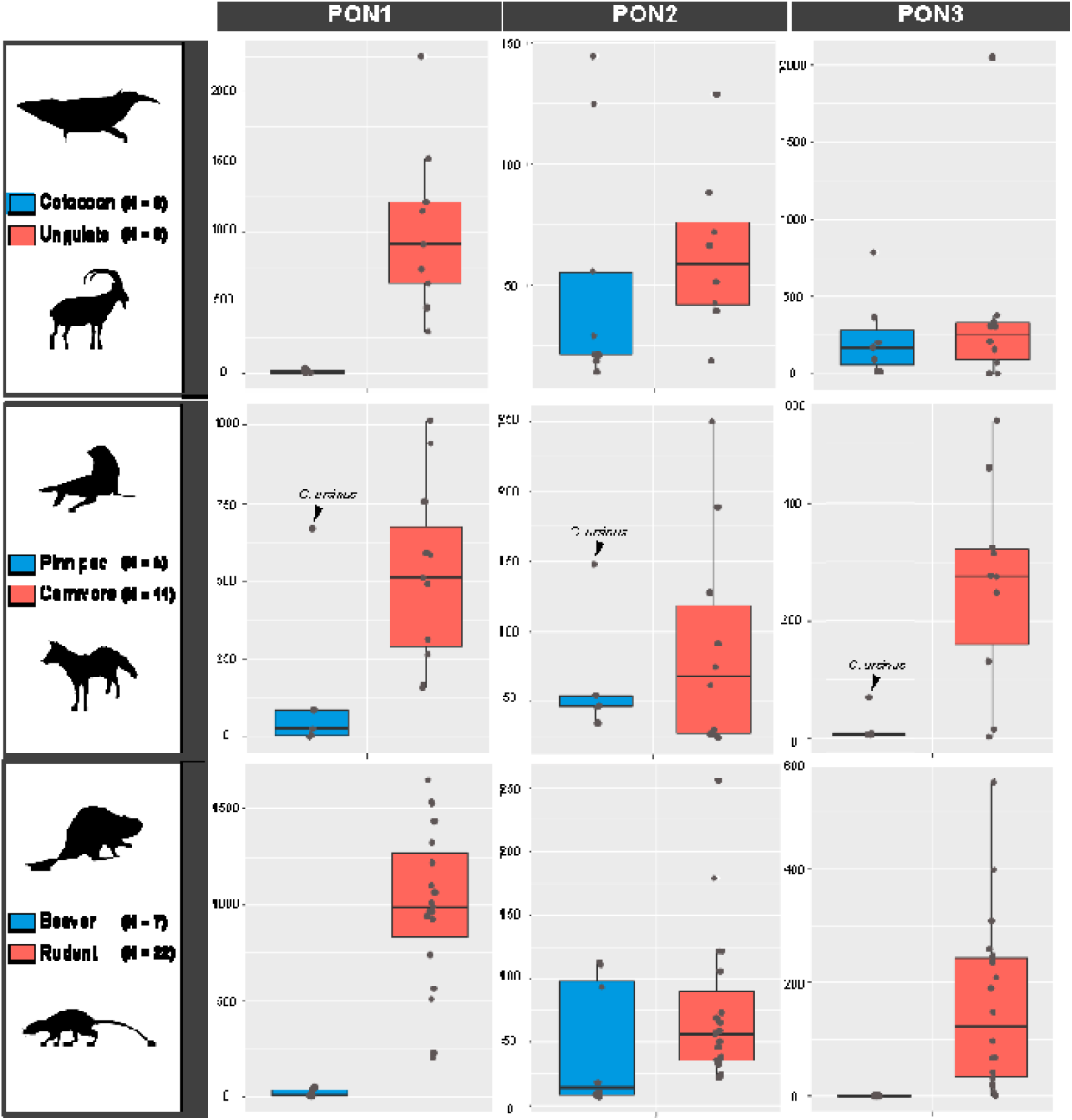
Transcriptional activity levels for the three *Pon* genes in three lineages with independent aquatic transitions (TPM; Transcripts per Million). Each dot represents a value for a singular species – some species had multiple individuals, and therefore their dot is an average value (Supplemental Table S1).

**Figure 4:**
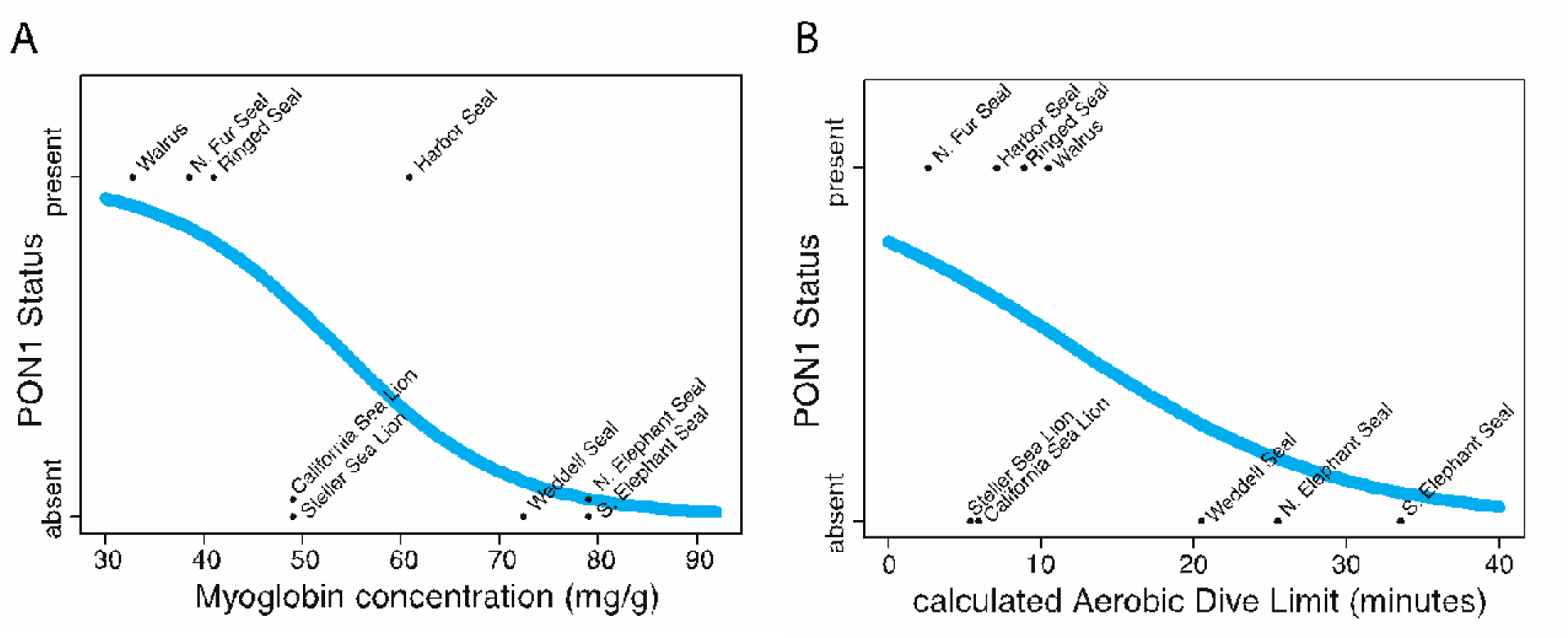
Fitted phylogenetic logistic regression of Pon1 status against (A) predicted myoglobin concentration in mg/g (*P* = 0.051) and (B) calculated aerobic dive limit in minutes (*P* = 0.1223), estimated using data for nine pinniped species. In (A), both California and Steller Sea Lion have no functional Pon1 and the same predicted myoglobin concentration.

### Evidence of Pon1 functional loss in some otter species

Another semiaquatic group in Carnivora is Lutrinae (otters). With DNA sequence available for eleven extant otter species (Figure 1), we predict loss of *Pon1* function from genetic lesions in the northern and southern sea otters (*Enhydra lutris* subsp.), which have diverged within the past 100 KYA (Beichman, et al. 2019). Ten of the eleven species (all except the North American river otter, *Lontra canadensis*) showed evidence of substitutions in *Pon1* at sites predicted to be functionally relevant from experimental studies (Supp Table S4); most of these sites are 100% conserved across non-aquatic mammals. The sea otter (*Enhydra lutris*) had substitutions at a conserved site that binds to catalytic calcium (N270), and this species and the giant otter (*Pteronura brasiliensis*) had substitutions at conserved active site wall residues (F222 and Y71, respectively). The Asian small clawed otter (*Aonyx cinereus*), spotted-necked otter (*Hydrictis maculicollis*), African clawless otter (*Aonyx capensis*), and sea otter had substitutions at conserved sites at which experimentally derived mutants were observed to have reduced activity against Pon1 substrates (H134, H155, W202, V304; Supp Table S4). Molecular evolutionary analyses supported elevated *d_N_/d_S_*(ω) in *Pon1* in the otter clade as a whole, with a point estimate of ω = 0.6579 across all otter lineages in contrast with a point estimate of ω = 0.2951 for all other mustelids. The elevation of ω in otters is consistent with relaxed selective constraint. A branch model with two different ω values for otters and other mustelids was a significantly better fit than a model with just one ω across all mustelids (LRT *p* = 0.010). Enzymatic results from two otter species sampled showed variable activity patterns, with the North American river otter having high activity closer to that of non-aquatic mustelids, while the Asian small-clawed otter had intermediate activity more similar to some of their semiaquatic carnivore relatives (*i.e.*, Odobenidae and Phocidae; Figure 2).

### Loss of Pon1 function and a surprising loss of Pon3 function in beavers

The American beaver, which is an adept diver, has an observed substitution in *Pon1* at amino acid H184, which is part of the Pon1 active site and was otherwise conserved across all other species for which sequences are available (Yeung, et al. 2008; Meyer, et al. 2018). *Pon1* coding sequences from additional semiaquatic rodent species, including one arvicoline rodent (muskrat) and one caviomorphid (capybara), showed only a few substitutions at functionally relevant sites. Both muskrat and capybara have a substitution at an amino acid that is part of the active site wall (R192), which is conserved in 85% of terrestrial lineages (including bats); muskrat showed an additional substitution at a site crucial for catalytic activity (I117), which is conserved in 94% of terrestrial lineages (Harel, et al. 2004).

Our comparison between beaver and terrestrial rodents involves multiple beaver liver RNA-seq datasets from 7 individuals compared to 19 different rodent species. Expression levels were significantly different for *Pon1* (*Z =4.009, P=6.1 x 10^-5^*) and *Pon3* (*Z =3.57, P= 0.0004*, but not for *Pon2* (*Z = 1.672, P= 0.095*) (Figure 3; Supp Table S2). Based on the beaver genome and transcriptome, *Pon3* is labeled as a pseudogene with no annotated expression (LOC109693470; *Castor canadensis* genome v1.0). We confirmed this by adding the *Pon3* CDS to the reference transcriptome and then mapping sequences, which ultimately recovered zero matches. This severe reduction in transcriptional activity is corroborated by the fact that the enzymatic activity against *Pon1* substrates in beavers is similar to that of the *Pon1* knock-out mouse (*Pon1*^-/-^), compared to that of their rodent relatives (Figure 2).

It is important to note that there is no transcriptomic data available for any semiaquatic rodent beyond the American beaver and the water vole. The expression of *Pon1* in the water vole is higher than any of the other rodent species assayed (1654 TPM). While beavers can dive for up to 15 minutes, the water vole is not known for spending any extended period of time diving; in addition, it is not truly adapted to the water, including a lack of webbed feet (Clausen and Ersland 1968).

### Loss of Pon1 function and expression in cetaceans

Pseudogenization of *Pon1* in cetaceans occurred ∼53 Mya, with all 5 representatives with sequenced genomes showing numerous genetic lesions, including multiple frameshifts and premature stop codons that are shared and must have occurred in the last common cetacean ancestor (Meyer, et al. 2018). Given this prior evidence, the expectation would be that expression would be null or very low. When comparing expression to ungulate relatives, there were significant differences found in *Pon1* expression (*P= 0.0001748*), but not for *Pon2* (*P= 0.189*), *Pon3* (*P= 0.613*) or *Actin* (*n1 = n2 = 7, P* = *0.053*) (Figure 3; Supp Table S2). The cetacean results serve as a control since their loss of *Pon1* function is ancient, and interestingly there is still detectable transcript, albeit at very low levels (0 - 26 TPM).

### Pon1 expression levels show moderate correlation with enzymatic activity

All semiaquatic species for which we analyzed data show reduced *Pon1* expression compared to their non-aquatic relatives, regardless of the presence or absence of genetic lesions. Within the species for which we have both expression and enzyme activity, there are five semiaquatic and aquatic species, including two pinnipeds (*Zalophus californianus, Phoca largha*), a rodent (*Castor canadensis*), and one cetacean (*Tursiops truncatus*) (Figure 1). The rest are non-aquatic lineages, including two rodents (*Mus musculus, Rattus norvegicus*), three carnivores (*Canis lupis familiaris, Felis catus, Mustela furo*), and three ungulates (*Ova aries*, *Sus scrofa*, *Capra hircus*). Comparing chlorpyrifos oxonase (cpo) activity levels against log TPM counts for *Pon1* showed a moderately strong relationship (R^2^= 0.758).

### Differentially expressed antioxidant genes tend to have lower expression in semiaquatic compared to non-aquatic lineages

After matching gene identity across references, a total of 108 antioxidant genes were assessed for expression differences in each of the (semi)aquatic/non-aquatic lineage pairs (Supp Table S5). Within beavers-rodents, we were able to compare 84 of those genes, with 34 showing significantly different expression patterns (40.4%). A total of 33 of the 34 genes were all marked by decreased expression in beavers compared to rodents, with only 1 gene (*MGST2*) showing increased expression. For cetaceans-ungulates, 86 antioxidant genes were compared, with only 26 (30.2%) showing significant differences in expression. Of those, 18 had decreased expression while 8 showed increased expression levels in cetaceans, including *APOA1*, *APOA4*, *GPX3*, *LTC4S*, *PRDX1*, *PRDX2*, *PRDX4*, and *NXNL1*. For pinnipeds-carnivores, 87 antioxidant genes were compared, of which 13 showed significant expression differences (14.9%); ten of those genes showed a decrease in expression in pinnipeds, while three (*APOA1*, *APOA4*, *GGT1*) showed increased levels of expression (Supp Table S5).

### Pon1 loss of function correlates best with diving-related phenotypes

Because Pon1 is thought to mitigate oxidative damage to bloodstream lipids, we tested for an association between loss of function and six phenotypes that differ between aquatic and terrestrial mammals and could also impact bloodstream lipid oxidation (Mackness, et al. 1991; Rosenblat and Aviram 2009). The first three phenotypes were related to dietary fatty acids because omega-3 and omega-6 fatty acids have different capacities to sustain oxidative damage (Miyashita, et al. 1993) and marine and terrestrial diets are known to have differing ratios of omega-3 to omega-6 fatty acids (Koussoroplis, et al. 2008), with omega-3 being more prevalent in marine ecosystems and more resistant to oxidation. Dietary phenotypes examined were (1) total omega-3 in diet, (2) total omega-6, and (3) omega-6:omega-3 ratio. Additionally, because large amounts of oxidative damage are accumulated while diving (Cantú-Medellín, et al. 2011; Vázquez-Medina, et al. 2012), we also explored the possibility that increased oxidative stress due to diving could be related to *Pon1* loss. We tested for an association between *Pon1* functional loss and (4) calculated aerobic dive limit (cADL), (5) maximum reported dive time, and (6) estimated myoglobin concentration. We used previously estimated cADL, the maximum amount of time an animal is predicted to be able to dive while subsisting on aerobic respiration (Costa, et al. 2001). Myoglobin concentration, as estimated from myoglobin surface charge, was previously shown to correlate well with current and ancestral diving ability across aquatic and semiaquatic mammals (Mirceta, et al. 2013).

For each phenotype we used a phylogenetic logistic regression to test its correlation with a binary *Pon1* functional status variable, encoded as present or absent, as judged from all available data including lesions, expression level, and enzymatic activity (details in Methods). None of the tests reached statistical significance, but the best predictor was myoglobin concentration with an uncorrected P-value of 0.051 (Figure 4; Supp Table S6). Over all tests, the diving-related phenotypes predicted *Pon1* functional status better than the dietary phenotypes; diving-related model fits had consistently lower Akaike Information Criterion (AIC) values than those for diet (maximum diving-related AIC 14.83; minimum diet-related AIC 16.41; Supp Table S6). In fact, the dietary variables correlated with *Pon1* status in the opposite direction from that hypothesized; for example, the estimated coefficient for total omega-3 (the best fitting diet variable; Supp Table S6) was 38.9, indicating that more omega-3 fatty acids would predict an increased probability of functional *Pon1*. If diet were the driving force behind *Pon1* functional loss, an increase in marine-prevalent omega-3 fatty acids would predict a decreased probability of functional *Pon1*. The lack of significance for these tests may be due to the small sample size (n = 9 species), since diving data are unavailable for most pinniped species.

## DISCUSSION

Previous work has shown that multiple clades of exclusively aquatic (cetaceans, sirenians) and semiaquatic lineages (pinnipeds, castorids, lutrids) – have each independently lost the function of the Pon1 protein at least once (Meyer, et al. 2018). Whereas the functional losses in cetaceans and sirenians were ancient, evidence suggests that 2 independent functional losses occurred in pinnipeds more recently, with some extant species retaining at least partial function of the Pon1 enzyme. With new evidence presented here from several clades, we estimate there to have been at least six independent losses of *Pon1* across mammals, all of which occurred in aquatic lineages. Given the importance of *Pon1* function for protection against toxicity from organophosphate byproducts (Furlong 2007), which may be present due to agricultural runoff in semiaquatic species’ habitats, it is critically important to characterize the extent of *Pon1* loss in all aquatic species. Gaining a more complete picture of *Pon1* functional loss throughout the mammalian phylogeny also enables us to test hypotheses regarding the specific selective pressures that may have led to this loss (Meyer, et al. 2018).

### Evolution of Pon1 and Pon3 Function

Our results here demonstrate that loss of *Pon1* function accompanies every mammalian transition to aquatic lifestyle that we have been able to assess, with one exception, the American mink. The six functional losses we observed apparently happened rapidly in most cases, with high inferred *d_N_/d_S_*implying that *Pon1* had not been subject to selective constraint on the majority of each ancestral branch of extant aquatic or semiaquatic species lacking *Pon1* function. In contrast, *Pon1* function has been highly conserved among terrestrial mammals, with terrestrial carnivorans, rodents, and ungulates retaining high levels of blood plasma activity against Pon1 substrates (Meyer, et al. 2018) (Figure 2). We also report here for the first time that *Pon3* has similarly evolved very low expression levels specifically in semiaquatic species, the beaver and pinnipeds. This striking co-evolution of *Pon1* and *Pon3* function with aquatic transition provides strong evidence that some characteristic of aquatic lifestyle removes purifying selection on *Pon* function or even favors its loss.

Loss of protein function is generally thought to be detrimental and therefore selectively disadvantageous; however, if a nonessential gene is pseudogenized, there may be no selective disadvantage and such pseudogenization can be selectively neutral. Recent work has suggested that gene loss may serve as an engine of evolutionary change (Olson 1999a), and gene loss of function has been implicated in numerous lineage-specific traits presumably crucial for fitness, across a variety of taxa (Zhao, et al. 2010; Meredith, et al. 2011; Yohe, et al. 2017; Esfeld, et al. 2018; Graham and Barreto 2020). Empirical studies have primarily characterized pseudogenes by identifying substitutions in coding regions, such as premature stop codons, frameshift mutations, and nonfunctional splice-sites; once such a nonfunctional mutation has fixed, pseudogenes are expected to evolve without selective constraint and to degenerate quickly. However, losses of function can also be caused by the disruption of promoters or other regulatory regions, reducing gene expression (Force, et al. 1999; Lynch and Force 2000). This phenomenon (*i.e.*, promoter degradation) has been extensively studied in plants, is largely facilitated by the promoter having few subfunctions, and is attributed to transposable-element insertion events (Woodhouse, et al. 2010; Yang, et al. 2011; Ochi, et al. 2017).

### Pseudogenization of Pon1 is preceded by reduction in transcriptional activity in multiple aquatic/semiaquatic lineages

Variation in gene regulation has been shown to underlie a significant proportion of phenotypic variance among closely related organisms (Levine and Tjian 2003) and is a considerable source of variation upon which natural selection can act. If the loss of *Pon1* in aquatic lineages is driven predominantly by directional selection on gene expression levels, and not drift, the expectation is that optimal expression in aquatic or semiaquatic species would shift relative to that of their terrestrial relatives. However, if stabilizing selection on gene expression is relaxed in aquatic or semiaquatic mammals, and the functional loss in these species is facilitated mostly through drift, the expectation would be an increase in expression variance within and among these species (Stern and Crandall 2018).

Ultimately, our results show a consistent, significant depression of *Pon1* transcriptional activity in aquatic and semiaquatic lineages (Figure 3; Figure 5 – Step 2), followed or accompanied by pseudogenization via accumulation of changes to coding sequence (Figure 5 – Step 3). This pattern is consistent with either selection for reduced *Pon1* expression in aquatic environments, or relaxation of selective pressures maintaining high levels of expression.

**Figure 5:**
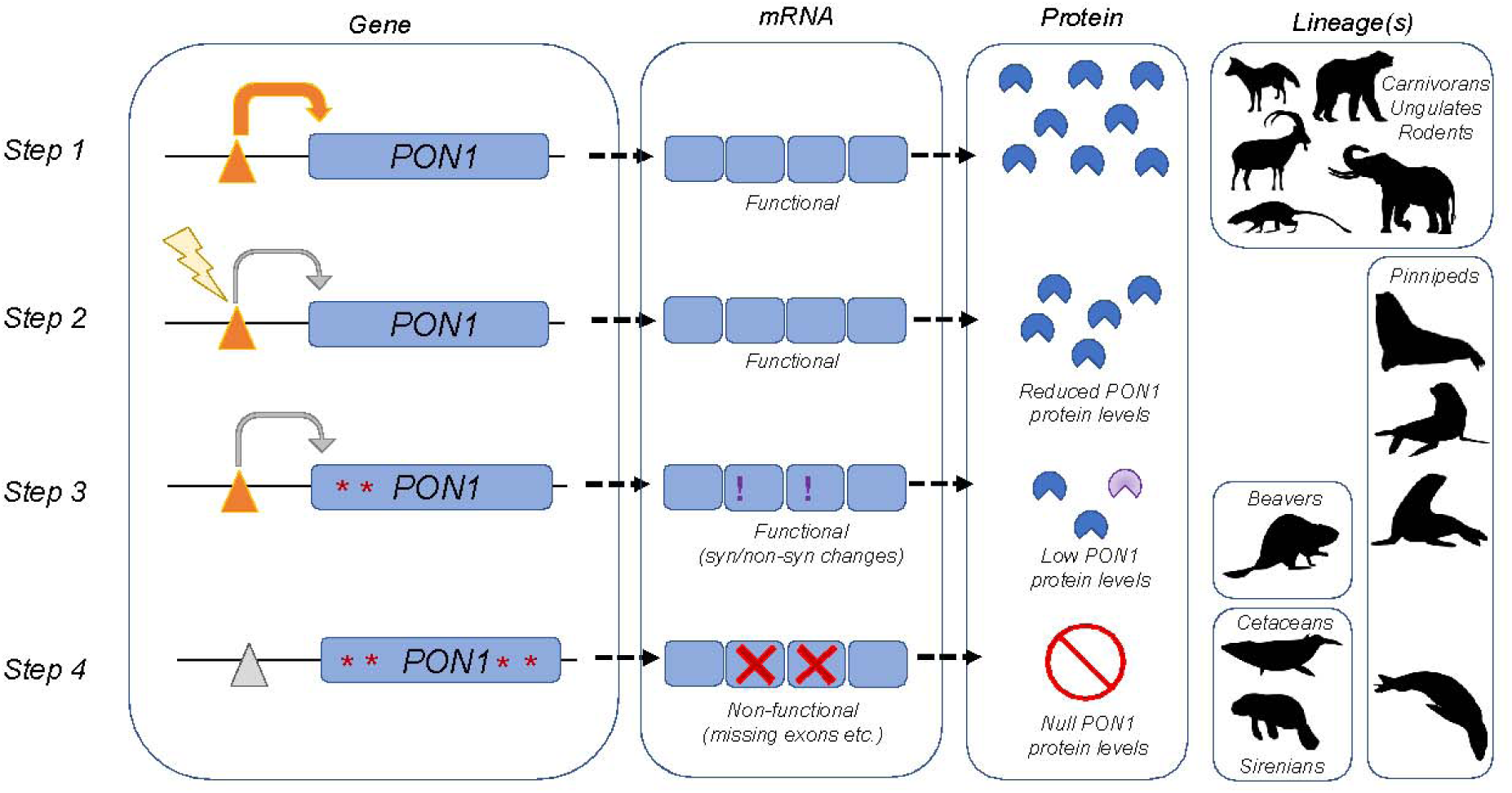
Schematic showing the inferred process by which Pon1 loses function during adaptation to aquatic environments. In Step 1, Pon1 is fully functional. In Step 2, a change to a regulatory element reduces transcription and hence Pon1 protein levels. In Step 3, the coding sequence of *Pon1* acquires further changes influencing its structure and/or function, leading to low and potentially dysfunctional protein. In Step 4, Pon1 function is fully lost, through a combination of changes preventing transcription and nonsense changes to the coding sequence (lesions), leading to no detectable RNA or protein. Silhouette images are from Phylopic (CC BY_SA 3.0).

Specifically, this same pattern of transcriptional reduction and accumulation of coding sequence substitutions impacting function occurred at least 4 separate times in aquatic or semiaquatic lineages, independently in cetaceans and beavers, and twice in pinnipeds (Figure 1; Figure 3). Expression reduction could also have occurred in sirenians and otters, but we were not able to get liver tissue from those species. It is currently unknown what amount of expression is required to meet levels necessary for “normal/wild type” *Pon1* function, although results from non-aquatic linages with an aquatic sister lineage show average expression levels ranging from 73 – 2251 TPM, compared to aquatic or semiaquatic levels of 0.1 – 62 TPM (Supp Table S2).

It is often assumed genes with any deactivating changes to their coding and promoter regions are “dead” (Balakirev and Ayala 2003), but a substantial number of disrupted genes are still actively expressed (Yamada, et al. 2003; Cheetham, et al. 2020). In our study, even in species in which *Pon1* is clearly a pseudogene, pseudogenization has not caused a complete loss of *Pon1* mRNA production. For example, Pon1 enzyme activity is effectively null in cetaceans, and *Pon1* was shown to have pseudogenized in this clade roughly 53 Mya, yet non-zero levels of transcript still persist.

### Requirements for extreme diving ability could cause Pon1 loss in aquatic mammals

Understanding the underlying physiological reasons facilitating pseudogenization of *Pon1* would allow us to better understand whether the repeated loss is truly adaptive, or due to drift. Meyer, et al. Meyer, et al. (2018) initially suggested that *Pon1* functional loss could be due to its protein’s role in either diving or diet; this study includes more species as well as transcriptional activity to better test these specific hypotheses.

Based on our analyses, the hypothesis that diving and oxidative stress impact *Pon1* loss in pinnipeds was better supported than the hypothesis that *Pon1* functional loss occurred due to an increased amount of omega-3 fatty acids in the aquatic diet, though neither set of tests was statistically significant (Figure 4; Supp Table S6). There are several factors influencing the power of these analyses - first, because of our stringent criteria for what qualified as *Pon1* loss of function (DNA lesions, no plasma activity, or RNA TPM ≤ 41.3) and functional (plasma activity or RNA TPM ≥ 61.76), our sample size for these analyses was 9 species. Second, for many of the species in this study, the loss of *Pon1* occurred on ancestral branches in animals whose phenotypes may not match their modern-day counterparts. For example, the ancestor of the California sea lion and the Steller sea lion lost *Pon1* function, but the sea lions had an explosion of diversity in the Miocene, with most members of the clade going extinct by the modern era (Debey and Pyenson 2013). It is therefore possible that these animals had different diving or dietary traits than their modern counterparts.

Myoglobin (Mb) concentration was most predictive of *Pon1* loss. Like its distantly related protein hemoglobin, Mb is an iron-and oxygen-binding protein found in both cardiac and skeletal muscle of vertebrates. In general, Mb has a higher oxygen binding affinity than hemoglobin, and is involved in the storage and diffusion of oxygen within muscle (Gros, et al. 2010). Deep-diving marine mammals are known to have more stable Mb than that of non-diving mammals, and these divers’ Mb is also expressed at higher levels in cardiac muscle (Scott, et al. 2000; Samuel, et al. 2015; Isogai, et al. 2018). In addition, the Mb net charge is higher in diving than non-diving mammals, potentially representing adaptation to prevent Mb self-association when it is at the high concentrations necessary to increase oxygen storage in muscle (Mirceta, et al. 2013). Maximal active dive time has been linked to the maximal skeletal muscle concentration of Mb (Kooyman and Ponganis 1998; Noren and Williams 2000; Lestyk, et al. 2009; Ponganis, et al. 2011). Deep diving mammals rely on oxygen stores in the blood and the muscle (80-90% of O_2_ storage), while shallow divers rely more on respiratory stores (Kooyman and Ponganis 1998).

Previous work has suggested increased Pon1 protects against atherosclerosis and lipid oxidation in a dose-dependent manner (Shih, et al. 1998; Tward, et al. 2002). Some human epidemiological studies have revealed an association between low Pon1 levels and an increased risk for coronary artery disease (Mackness, et al. 1999; James, et al. 2000; Jarvik, et al. 2000). In contrast, meta-analyses across a large number of such studies have shown only a marginal relationship (Wheeler, et al. 2004), though that may be due to the distribution of the *Pon1* polymorphisms varying with ancestry (Davis, et al. 2009; McDaniel, et al. 2014). If a major function provided by *Pon1* is to protect against atherosclerosis, potentially by breaking down oxidized lipids, it is a paradox that these diving species would lose the function of this gene. Why would species that encounter more oxidative stress lose the ability to produce a functional enzyme whose presence is protective in that context? These evolutionary losses of *Pon1* and *Pon3* raise the possibility that their functional roles or the oxidative conditions induced by diving are not fully understood.

### Potential for inflammation to drive Pon1 and Pon3 functional loss across multiple lineages

It is important to note that diving involves a large set of physiological responses, one or more of which may be the causative factor(s) in driving repeated pseudogenization of *Pon1*. For example, prolonged diving typically involves organisms drawing down their body oxygen stores, which results in generalized hypoxemia and local tissue hypoperfusion and hypoxia (McDonald and Ponganis 2013). Such extended bouts of hypoxia incur cytokine-mediated inflammation; however, animals that are adapted to hypoxic environments (high-altitude) or engage in hypoxic behaviors (i.e., diving, burrowing) are thought to have a largely blunted inflammation response or to have adopted additional mechanisms to deal with hypoxia-induced chronic low-grade systemic inflammation (Beall, et al. 2002; Erzurum, et al. 2007; Okin and Medzhitov 2012; Zhang, et al. 2016). This is because hyperactivation of the inflammatory response is detrimental, potentially causing systemic inflammatory response resulting in septic shock, as well as causing cardiovascular disease, among other negative consequences (Libby 2006; Medzhitov 2008).

Pon1 activity has been shown to have antioxidant and anti-inflammatory properties (Ng, et al. 2008; Marsillach, et al. 2009; Mackness and Mackness 2010; Aharoni, et al. 2013); thus, the pseudogenization of *Pon1* would at first glance seem counter-intuitive, as inactive Pon1 would not be able to suppress the inflammatory response. However, it is possible that additional mechanisms have counteracted the predicted increase in inflammation in diving mammals due to hypoxia. For example, Tibetan and Andean human populations show no sign of systemic inflammation compared to low-landers, and studies have shown evidence of multiple mechanisms that aid in circumventing hypoxia, as well as the resulting inflammation, in these populations; these include changes to the Nitric-Oxide pathway (Erzurum, et al. 2007), TNF-mediated pathways (Pham, et al. 2021), and the HIF-pathway. Perhaps *Pon1* function is lost in aquatic species because they have similarly developed reduced inflammation and protection from oxidative stress by other means. On the other hand, Pon1 and Pon3 were shown to hydrolyze eicosanoids, which are lipids that can modulate inflammatory and immune responses (Teiber, et al. 2018). *Pon1* functional loss in aquatic species could have occurred due to needed changes in eicosanoid signaling to reduce inflammation.

Inflammation is a difficult phenotype to measure since there is no easy way to quantify its activity, especially during such a transient behavior as diving; instead, we sought to assess Pon1 activity in other lineages which have [a] evidence of accelerated evolution of *Pon1* and [b] a unique inflammatory response (*i.e.*, bats). Previous work showed similar acceleration of Pon1 evolutionary rate in bats to that of marine species; in addition, although bats did not show evidence of deterioration of the *Pon1* gene, some bats had distinct substitutions which potentially would affect activity (Meyer, et al. 2018). Recent work in bats has shown they have a dampened inflammatory pathway – specifically, bats have multiple strategies that reduce proinflammatory responses, thereby mitigating potential immune-mediated tissue damage and disease (Ahn, et al. 2019; Goh, et al. 2020; Larson, et al. 2021). In addition, there is evidence for loss of an entire gene family *PYHIN* (Ahn, et al. 2016), as well as suppressed expression of *c-Rel* (Larson, et al. 2021), both of which have known roles in inflammasome sensing, in bats. Our results show that chiropterans also have a pronounced reduction in *Pon1* expression (median 362 TPM) compared to closely related lineages, which include carnivores (median 480 TPM; Mann-Whitney U test P>0.05), ungulates (median 914 TPM; P<0.05), or both combined (median 613 TPM; P<0.05) (Supp Table S2).

Reactive oxygen species (ROS) are frequently linked to initiating the inflammatory response; thus, we might expect the deficit of serum paraoxonase activity to be compensated for by an increase in expression of other antioxidant genes in semiaquatic species. However, our comparison of expression patterns of antioxidant genes in aquatic and semiaquatic lineages to those in their non-aquatic relatives yielded evidence of a general decrease in antioxidant expression in aquatic species relative to non-aquatic species (Supp Table S5). This may suggest that semi-aquatic species have additional mechanisms to prevent the initiation of the inflammatory response by ROS, paralleling the complex inflammation prevention mechanisms that accompany substantial changes in Pon1 in bats. Taken together, the reduced expression of Pon1 in bats and the reduced expression of antioxidant genes in semiaquatic species lend some credence to the notion that circumventing an otherwise deleterious inflammation response may be the driving factor behind changes in *Pon1* evolutionary rate (bats, pinnipeds) and subsequent pseudogenization (pinnipeds).

## ACKNOWLEDGEMENTS

We would like to thank Maria Chikina, Johanna Kowalko, and members of the Clark and Meyer labs for helpful discussions, as well as anonymous reviewers for their constructive suggestions to improve the manuscript. We would like to thank the University of Alaska – Museum of the North, the Burke Museum – University of Washington, and the Utah Museum of Natural History for granting us access to the tissue samples we requested. In addition, we would like to thank the otters of the Loveland Living Planet Aquarium for granting us access to their blood serum. Blood and plasma samples for Steller sea lions were collected under NOAA permit no. 14326. Any use of trade, firm, or product names is for descriptive purposes only and does not imply endorsement by the U.S. Government.

This work was supported by a NIH T32 Hematology Fellowship and NIH K99/R00 Pathway to Independence Fellowship to AMG (NIH 3T32DK007115); a UPMC Hillman Cancer Center Academy Summer Internship to AM; a Genomics Summer Research for Minorities internship to MB (NIH 5R25HG009886); and R01 HG009299 and R01 EY030546 NIH grants to NC.

## Data and code availability

Sequence from samples generated in this study are available on the NCBI (PRJNA805216), and additional code and data are available in a Dryad repository (https://doi.org/10.5061/dryad.612jm6452).

Any additional information required to reanalyze the data reported in this paper is available from the lead contact upon request.

## MATERIALS AND METHODS

### Sample Overview

Tissue samples from a total of 30 individuals across 12 species were obtained through the University of Alaska – Museum of the North and through Burke Museum of Natural History and Culture - University of Washington (Supp Table S7), via the ARCTOS collaborative collection management website. These samples were subsequently used for Pon1 sequencing and for RNA-seq analysis. We obtained blood samples for DNA sequencing and Pon1 enzyme activity testing for mink from a mink farm in Ohio, for harbor seals from Six Flags Discovery Kingdom in California, for Steller sea lions from NOAA and the University of Alaska Fairbanks (NOAA permit no. 14326), for northern elephant seals from Sonoma State University (NMFS permit # 19108), and for several otter species from the Loveland Living Planet Aquarium in Utah. Blood samples from all animals in human care were obtained using procedures approved by the University of Utah and each sampling institution’s research review committees prior to conduction. Additional Pon1 CDS sequence information was acquired from publicly available genomes (Supp Table S8) and transcriptomes (Supp Table S9). Expression data in liver samples for aquatic and semiaquatic mammal lineages and their close non-aquatic relatives were obtained through NCBI Short Read Archive (SRA), for a total of 90 individuals from 56 different species across Carnivores (14 species), which include Pinnipeds (2), plus Ungulates (10), Cetaceans (17), and Rodents (20), which includes Beaver (Supp Table S9).

### Determining Pon1 and PON3 Sequence from Public Genomes

A combination of BLAT, BLAST, bowtie2, and MAKER annotation were used to find the Pon1 coding sequence within publicly available genomes of new species of carnivores, and rodents. Specifically, for the Steller sea lion and northern fur seal, in BLAT (Kent 2002) the first and last exon of the walrus Pon1 sequence were used as queries against the entire genomes with default parameters; the coordinates from the BLAT searches were then input into a script which then extracted the sequences between the first and last exons, inclusively, from each species. The Pon1exons in these regions were then further identified based on homology to the walrus sequence. Finally, the exons were manually checked for any lesions or substitutions at important sites, using walrus (*Odobenus rosmarus divergens*) as a reference.

For mink, California sea lion, capybara, giant river otter, and nutria genomes we used BLAST to identify the Pon1 exons. Each exon of Pon1 from the closest relative available for each species (ferret for mink, sea otter for giant river otter, Steller sea lion for California sea lion, and guinea pig for capybara + nutria) was used as a query against the relevant species’ genome. If nucleotide BLAST returned no results for an exon, we then translated the sequence and subjected the translation to a tBLASTn search against the genome. If the exon still showed no results, we then used the region containing the exon plus some surrounding sequence (10 bp) as a query in a BLAST against the genome. The alignment coordinates from the BLAST searches were then used to extract the sequences of each exon plus approximately five base pairs on either side from the genome. Splice sites were then identified, exons manually concatenated and then translated. These sequences were then aligned separately with other carnivorans and rodents, using Seaview v4.6.1 (Gouy, et al. 2010), where they were manually checked for lesions and/or substitutions at potentially important amino acid sites, using dog (*Canis lupis familiaris*) as a reference for Carnivora and mouse (*Mus musculus*) as a reference for Rodentia.

For six otter species, the whole-genome sequence was downloaded from NCBI SRA (PRJNA841998) and sequences mapped to the *L. canadensis* full gene Pon1 reference using bowtie-2 (Langmead and Salzberg 2012). A consensus sequence was generated for each species, coding regions annotated, and then translated; these protein sequences were then aligned using MUSCLE to the human Pon1 (Uniprot P27169) which had important sites flagged (Supp Table S4).

### Gene Sequencing and Lesion Identification from Blood and (Museum) Tissue Samples

Sequencing was performed for samples from an otter (blood buffy coat), beaver (whole blood) and pinnipeds (liver, blood, muscle). Both RNA and DNA were extracted from any samples with liver tissue - samples were ground in 1.5 mL tubes using a Bel-Art™ SP Scienceware™ Liquid Nitrogen-Cooled Mini Mortar, and then DNA and RNA was extracted using the Trizol protocol. For the blood and muscle tissue samples, DNA was extracted using the Qiagen DNeasy Blood and Tissue kit. For pinnipeds, reference sequences of Pon1 were acquired from 6 different pinniped species with genomes available, including *Zalophus californianus* (California sea lion; NC_045606), *Neomonarchus schauinslandi* (Hawaiian monk seal; NW_018734323), *Callorhinus ursinus* (Northern fur seal; NW_020312848), *Eumetopias jubatus* (Steller sea lion; NW_020998611), *Phoca vitulina* (Harbor seal; NW_022589707), and *Halichoerus grypus* (Gray seal; NW_023400273). For otters, reference sequences of Pon1 were acquired from 3 different otter species with genomes available, including *Lontra canadensis, Lutra lutra and Enhudra lutris kenyoni.* For beaver, reference sequences of Pon1 were acquired from 4 different *Castor canadensis* genomes available on NCBI (C.can genome v1.0; ASM982264v1; CasCan_v1_BIUU; bgp_v1).

For each group, the whole gene region was aligned using MUSCLE (Edgar 2004) and then degenerate primers were created for each of the 9 exons using Primer3 (Untergasser, et al. 2012) (Supp Table S10). Using the DNA extracted from museum samples, each exon was subjected to PCR (NEB OneTaq®, M0480) and sequencing (University of Utah; HSC Core Facilities). All 9 exons were then combined into complete CDS and compared to the consensus CDS generated from the genome references. Potential genetic lesions, substitutions, as well as conservation of the 10 major active sites, were identified using criteria described in Meyer, et al. (2018), facilitated by the Geneious Prime software (Kearse, et al. 2012). The PCR products were sequenced by the University of Utah Genomic Research Core via Sanger sequencing.

### Expression Analysis of Pon1 in liver derived RNA-seq

Expression patterns using transcriptomic data was performed for RNA-seq libraries created from museum liver samples, as well as liver RNA-seq libraries available on the Short Read Archive (SRA; Supp Table S7, S9).

For museum samples, the extracted RNA for 4 samples was sent to Novogene for library preparation (polyA enrichment) and sequencing via NovaSeq PE150. An additional 13 individuals had RNA-seq libraries made using Swift Biosciences RNA library prep kit, before being sent to Novogene for sequencing – these libraries were trimmed for adapters and quality filtered via BBduk in the BBtools package (Bushnell 2018). However, 4 libraries/individuals were used in further analyses due to clear RNA degradation (RNA Integrity Number = RIN < 4 ; SUPP Table S7). Data downloaded from the SRA included transcriptomic data derived exclusively from liver for other pinniped species, as well as other carnivores, cetaceans, ungulates, rodents and chiropterans (Supp Table S9). Each library was mapped using Salmon (Patro, et al. 2017) against the closest-related reference available – for pinnipeds, they were mapped to all 6 pinniped references available. When possible, multiple individuals were used for the same species to assess within-species expression variation. Regardless of origin, expression levels of Pon1, Pon2, Pon3, and multiple endogenous control genes were assessed in all transcriptomes (Supp Table S2, S11).

For all samples run through the Salmon pipeline, differences in expression between aquatic or semiaquatic and non-aquatic lineages for 109 antioxidant genes (Carbon, et al. 2009; Vázquez-Medina, et al. 2012) were tested for significance using a Mann-Whitney U. There were a total of 43 different transcriptome references used, thus each transcript was renamed according to the closest BLAST hit from the human transcriptome (GCF_000001405.39_GRCh38.p13_rna), and then isoforms for each gene collapsed into one per gene.

### Assays for Enzymatic Activity Against Organophosphate-Derived Substrates

Blood was collected in lithium heparin tubes and centrifuged at 1,500 - 10,000 x g for 10 - 15 min at 4 °C. Plasma was separated from the blood cell fraction and kept stored at −80 °C until use. All activity assays were determined in a SPECTRAmax® PLUS Microplate Spectrophotometer (Molecular Devices, Sunnyvale, CA). The assay values were corrected for path-length using the software SoftMax Pro 5.4 (Molecular Devices). Levels of plasma arylesterase (AREase), chlorpyrifos-oxonase (CPOase), diazoxonase (DZOase) and paraoxonase (POase) activities were determined kinetically as previously described (Richter, et al. 2009). Briefly, plasma of all the species analyzed were diluted 1/10 in dilution buffer (9 mM Tris-HCl pH 8.0, 0.9 mM CaCl_2_) and assayed in triplicate at either 37 °C (for CPOase and POase) or at room temperature (for AREase and DZOase). Activities were expressed in U/mL (AREase, CPOase, and DZOase) or in U/L (POase), based on the molar extinction coefficients of 1.31 mM-1 cm-1 for phenol (the hydrolysis product of phenyl acetate, AREase activity); 5.56 mM-1 cm-1 for 3,5,6-trichloropyridinol (the hydrolysis product of CPO); 3 mM-1 cm-1 for 2-isopropyl-4-methyl-6-hydroxypyrimidine (the hydrolysis product of DZO); or 18 mM-1 cm-1 for p-nitrophenol (the hydrolysis product of PO). Alkaline phosphatase was assayed in undiluted plasma of all species at 37 °C in triplicate as follows: The plasma sample, 10 µL, was added to 170 µL of 0.95 M diethanolamine pH 9.8, 0.5 mM MgCl_2_. The assay was initiated by adding 20 µL of 112 mM p nitrophenyl phosphate in water. The absorbance at 405 nm was followed for 4 min. Activities were expressed in U/L based on the molar extinction coefficient of 18.0 mM-1 cm-1 for p-nitrophenol (the hydrolysis product of p-nitrophenyl phosphate). Phenyl acetate (CAS 122 79-2, 99% purity), p-nitrophenyl phosphate (CAS 333338-18-4, ≥97% purity), and other reagent chemicals were purchased from Sigma-Aldrich. Chlorpyrifos oxon (CPO; CAS 5598-15-2; 98% purity), diazoxon (DZO; CAS 962-58-3; 99% purity) and paraoxon (PO; CAS 311-45-5, 99% purity) were purchased from Chem Service Inc. (West Chester, PA).

### Collection and calculation of Diving Metrics

We collected maximum dive times for seven pinniped species from Mirceta, et al. (2013), who gathered these data from multiple original studies (Slip, et al. 1994; Stewart and DeLong 1995; Teilmann, et al. 2000; Gjertz, et al. 2001; Plötz, et al. 2002; Sterling and Ream 2004; Eguchi and Harvey 2005) and for two species from Schreer, et al. (2001) (Supp Table S1). We obtained cADL (calculated aerobic dive limit) from literature if it was available (Schreer, et al. 2001; Weise and Costa 2007; Hassrick, et al. 2010; Weingartner, et al. 2012). For the Steller sea lion, cADL was not available, so we manually calculated cADL by dividing total body oxygen volume by metabolic rate (Gerlinsky, et al. 2013). If measurements for multiple individuals were available for cADL, we took the average of the values (Supp Table S1). Full data, references and results are available in repository folder “Diving_Variables”.

### Calculation of Diet Metrics

We took diet composition across 7 different food categories (Pauly, et al. 1998), with the fatty acid content for each category calculated as follows – (1) for benthic invertebrates, omega-3 and omega-6 values were averaged across clam and scallop (Santhanam 2014), (2) for large zooplankton, total lipid amount for *L.vannamei* was multiplied by the fraction of omega-3 and omega-6 lipids for *E. superba* (Bottino 1975; Santhanam 2014), (3) for squids, average values for omega-3 and omega-6 were multiplied by overall percent fat (Zlatanos, et al. 2006), (4) for small pelagic fishes, the values for total fat composition for *E. fimbriate* were used (Santhanam 2014), (5) for mesopelagic fishes, the values were calculated from the average of mesopelagic fishes (ie. percent fatty acid times percent total lipid content) (Stowasser, et al. 2009), (6) for miscellaneous fishes, total percent fatty acid by percent of the fatty acid type was multiplied for *P. niger* (Santhanam 2014), and (7) finally, fatty acid content for higher vertebrates was determined by multiplying the fatty acid values and percent fat by weight (Golet and Irons 1999; Wold, et al. 2011). If the percent total omega-3 for any category was unavailable, we used the total amount of EPA and DHA instead. The total amount of omega-3, omega-6 and omega-6/(omega-3+omega-6) fat was calculated for each prey category using a custom script (Determine_lipid_content.R; Results in repository folder “Dietary_Lipids”). These values were then multiplied by the fraction of each prey category in a species’ diet and added (Pauly, et al. 1998) to determine the overall omega-6:omega-3 ratio.

### Testing Correlations between Pon1 Loss and Potential Predictive Variables

We conducted phylogenetic logistic regressions of diet and diving variables against Pon1 status using the phyloglm and phylolm functions from the phylolm package (v2.6.2) along with the ape package (v5.6-2) in R (v4.1.2) (Ives and Garland Jr 2010; Tung Ho and Ané 2014). We assessed model fit using the Akaike information criterion. The full script is available as PGLS_Analysis_Pinniped.R in data repository folder “PGLS”. The status of Pon1 was determined to be “present” (encoded as ‘1’) if the species showed any plasma activity or RNA levels above 10. Pon1 status was determined to be “absent” (encoded as ‘0’) if any DNA lesions or substitutions in amino acid sites critical for function were present. Additionally, Pon1 status was made to be 0 if no plasma activity was seen or RNA levels were below 5 TPM. Species for which only a DNA sequence with no lesions were present were excluded from this analysis, as it has been shown that even without genetic lesions, some pinniped species lack Pon1 plasma activity (Meyer, et al. 2018). The species tree (PGLS_Tree.txt) used for all of these analyses was adapted from Higdon, et al. (2007) with branch lengths corresponding to millions of years. All scripts, data, and results are available in repository folder “PGLS”.

### Examination of Sequence Evolution in Lutrinae

To test hypotheses about sequence evolution in otters, we used branch model 2 in PAML (Yang 1997). We tested one alternative hypothesis -- that otters show a different rate of Pon1 sequence evolution from other mustelids -- against two null hypotheses. We modeled this alternative hypothesis by clustering the otters and the other mustelids separately and allowing *d_N_/d_S_*to vary between the two groups. We then compared this to a model where otters were clustered with the other mustelids, but *d_N_/d_S_* for this group was restricted to the *d_N_/d_S_* for other mustelids estimated in the alternative model, to simulate *Pon1* in otters evolving at the same rate as in the other mustelids. Finally, we fit a model where otters were clustered with the other mustelids, but *d_N_/d_S_* was allowed to vary freely for this group, representing a scenario where any shift in rates within otters is incorporated into *d_N_/d_S_* estimation for all mustelids. The alternative model was then tested against each null model with a log likelihood test with df = 1. For all models, other major groups in Carnivora (canids, feliformes, etc.) were clustered and *d_N_/d_S_*was determined for each group as a whole. We also fit a model where *d_N_/d_S_*was determined freely for each branch (branch model 1) to examine general changes in the rate of evolution for *Pon1* in otters.

## SUPPLEMENTARY FIGURE

**Figure S1:**
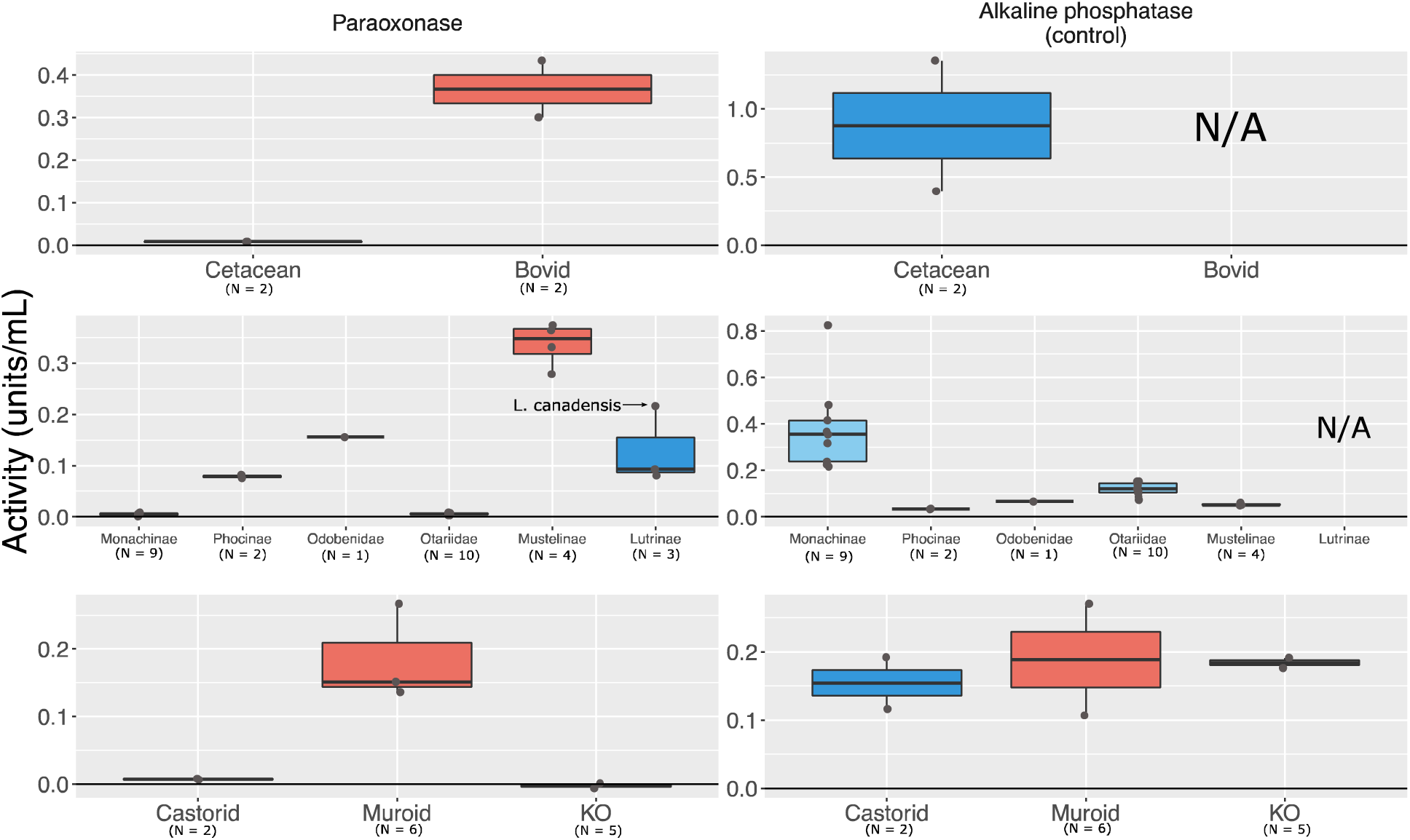
Blood plasma from semiaquatic species varies in activity against paraoxon, a traditional Pon1 substrate against which Pon1 is not protective for organophosphate toxicity. Levels of alkaline phosphatase activity are similar for aquatic, semiaquatic, and terrestrial species, serving as a control for the quality of the blood plasma tested. Plots show enzymatic activity in units/mL against paraoxon (L) and nitrophenyl phosphate (R) for aquatic/semiaquatic and closely related terrestrial species within (from top to bottom) Cetartiodactyla, Carnivora, and Rodentia. Carnivora species included are *M. angustirostris* (Monachinae), *P. vitulina* (Phocinae), *O. rosmarus* (Odobenidae), *E. jubatus* (9) and *Z. californianus* (1) (Otariidae), *M. furo* (Mustelinae), and *A. cinereus* (2) and *L. canadensis* (1) (Lutrinae). Data for Bovids (sheep, goat) and one Muroid (McGrath, et al.) are from (Furlong, et al. 2000b). See also Figure 2 and Table S12.

**Figure S2:**
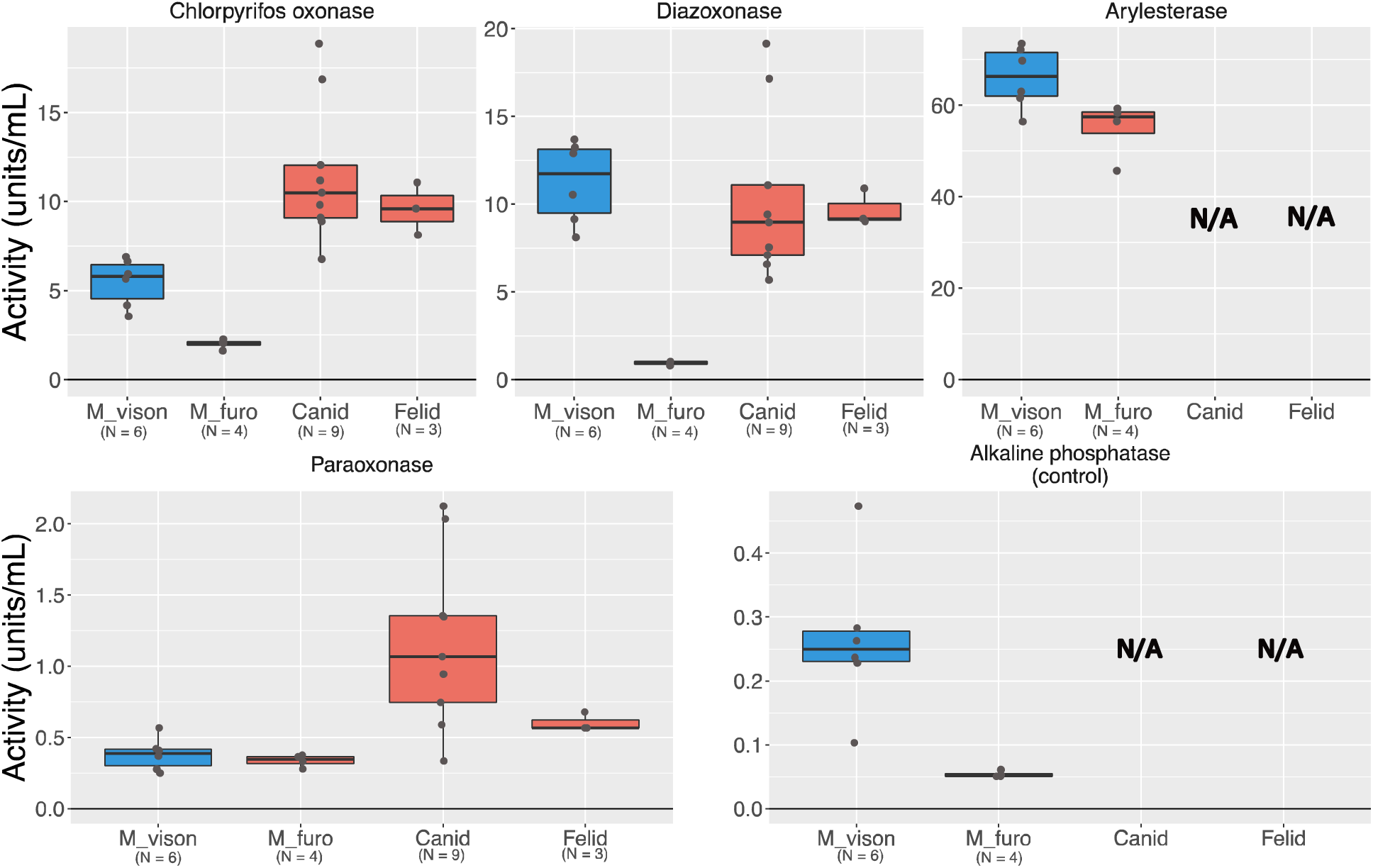
Blood plasma from the semiaquatic mink (*M. vison*) displays activity against Pon1 substrates that is comparable to that of terrestrial Carnivoran species. Plots show enzymatic activity in units/mL against chlorpyrifos oxon, diazoxon, phenyl acetate, paraoxon, and nitrophenyl phosphate (R) for *M. vison, M. furo, C. lupus familiaris,* and *F. catus*. Data for *C. lupus familiaris* and *F. catus* are from X.

